# A Statistical Model for Describing and Simulating Microbial Community Profiles

**DOI:** 10.1101/2021.03.26.437146

**Authors:** Siyuan Ma, Boyu Ren, Himel Mallick, Yo Sup Moon, Emma Schwager, Sagun Maharjan, Timothy L. Tickle, Yiren Lu, Rachel N. Carmody, Eric A. Franzosa, Lucas Janson, Curtis Huttenhower

## Abstract

Many methods have been developed for statistical analysis of microbial community profiles, but due to the complex nature of typical microbiome measurements (e.g. sparsity, zero-inflation, nonindependence, and compositionality) and of the associated underlying biology, it is difficult to compare or evaluate such methods within a single systematic framework. To address this challenge, we developed SparseDOSSA (Sparse Data Observations for the Simulation of Synthetic Abundances): a statistical model of microbial ecological population structure, which can be used to parameterize real-world microbial community profiles and to simulate new, realistic profiles of known structure for methods evaluation. Specifically, SparseDOSSA’s model captures marginal microbial feature abundances as a zero-inflated log-normal distribution, with additional model components for absolute cell counts and the sequence read generation process, microbe-microbe, and microbe-environment interactions. Together, these allow fully known covariance structure between synthetic features (i.e. “taxa”) or between features and “phenotypes” to be simulated for method benchmarking. Here, we demonstrate SparseDOSSA’s performance for 1) accurately modeling human-associated microbial population profiles; 2) generating synthetic communities with controlled population and ecological structures; 3) spiking-in true positive synthetic associations to benchmark analysis methods; and 4) recapitulating an end-to-end mouse microbiome feeding experiment. Together, these represent the most common analysis types in assessment of real microbial community environmental and epidemiological statistics, thus demonstrating SparseDOSSA’s utility as a general-purpose aid for modeling communities and evaluating quantitative methods. An open-source implementation is available at http://huttenhower.sph.harvard.edu/sparsedossa2.

## Introduction

Microbial community research has increasingly benefited from study designs inspired by molecular epidemiology, particularly with the goal of associating features of the human microbiome with health and disease [1]. This has enabled discoveries ranging from overall ecological dysbiosis in gut community structure during inflammatory bowel disease (IBD) [2] to specific microbial species, strains, and gene families linked to colorectal cancer (CRC) [3]. However, in almost all cases, existing statistical methods for genetic, transcriptional, metabolomic, or other molecular epidemiology cannot be accurately applied directly to microbiome measurements, due to their unique measurement error, noise, zero-inflation, compositional, and non-independence properties [4, 5]. This has led to inaccuracy issues in the literature, such as confounding, uncorrected population structure, batch effects, and a high rate of false positives [6–9]. There is thus an unmet need for statistical frameworks capable of capturing all aspects of microbiome epidemiology, both for the sake of accurately parameterizing and testing real community profiles, and for “reversing” parameterized models to simulate controlled, synthetic microbiomes for accurate methodology evaluation.

Transcriptional biomarker discovery has a similar history, in which early statistics to associate gene expression patterns with human phenotypes were met with challenges of noise, dimensionality, and test appropriateness [10]. This led to some of the first models for gene expression integrating features of underlying transcriptional biology, different assay platforms, and measurement noise [11]. These were in turn also “reversed” to provide simulated expression data for methods evaluation under guaranteed, controlled circumstances [12], permitting some of the first truly quantitative transcriptional epidemiology and comparative methods evaluation [13].

Models of microbial community structure are similarly important, and both their biological structure and measurement technologies are quite distinct from those for other sources of short-read sequence generation [14]. Microbial community profiles can be derived roughly equivalently from either amplicon (e.g. 16S rRNA gene) or metagenomic shotgun sequencing, and they consist of the (typically compositional) counts or proportions of taxa, genes, pathways, or other features derived from the source sequencing data. Like other types of molecular epidemiology profiles, they are typically a) high-dimensional (number of features equivalent to or surpassing sample size) [1] and b) require both feature-feature and sample-sample biological interactions (i.e. correlations or population structure) to be accounted for [15].

Additionally, microbiome data possess further unique properties that prohibit direct application of models from other molecular epidemiology research. They are considerably more sparse, i.e. zero-inflated, both due to low sequencing depth and biological absence [1]. As a result, in different settings, either biological presence/absence of microbial features or their abundances can be linked to phenotypes [16]. Microbial abundances from sequencing are also near-universally available only on a relative (compositional) scale, thus constrained to sum up to a constant. The combination of general high-dimensional statistical challenges with those unique to ecological profiles have impeded the development of a single, universal model of microbial feature structure.

As such, most previous strategies for modeling or simulating microbial community profiles (typically for methods evaluation) have been relatively simple [5]. Here, we will use “features” and “profiles” to refer to the quantification of taxa or other entities (e.g. genes or pathways) as counts or relative abundances from microbial community sequencing. McMurdie and Holmes [5] adopted deterministic mixing and multinomial sampling for simulating microbial taxa count observations; it thus does not allow for interaction between microbial features, nor does it model biological (as opposed to technical) absences. Similarly, Thorsen *et al.* [17] adopted random resampling of real-world data for simulating “new” microbial features and samples, indirectly violating compositionality and, again, excluding possible feature-feature interactions. Only recently, metaSPARSim [18] adopted a formal statistical model specifically for simulation of 16S rRNA gene amplicon-sequenced microbial observations (here abbreviated 16S), namely, the gamma-multivariate hypergeometric (gamma-MHG) distribution. However, the gamma-MHG model, itself an over-dispersed version of the multinomial model, still does not allow for biological absences or feature-feature interactions. Additionally, the model’s sampling implementation requires iteration over read depth for a given sample, which induces impractically high computation burdens to achieve realistic sequencing depths [1]. None of these frameworks formally capture microbial covariation with real or simulated covariates. In addition to other uses of such models, this is perhaps the most important aspect needed for benchmarking applications, where it enables estimation of power, false discovery rates, and effect sizes for microbiome epidemiology.

To address these gaps, we present SparseDOSSA (Sparse Data Observations for the Simulation of Synthetic Abundance), a statistical model that can be used to capture and, in turn, simulate realistic microbial community profiles. Motivated by the biological and technical data generation mechanisms and properties of microbial abundance observations, SparseDOSSA has model layers for a) zero-inflated marginal microbial abundances, b) penalized estimation of highdimensional feature-feature interactions, c) enforced normalization to address compositionality, and d) spiking-in of controlled microbe-microbe and microbe-environment covariation for benchmarking. We demonstrate through validations that the current implementation version, SparseDOSSA 2, accurately captures microbial community population and ecological structures across different environments, host phenotypes, and sequencing technologies, and is capable of recapitulating comparable, realistic synthetic profiles. We also show example applications in microbiome study design power analysis and in recapitulating a complex end-to-end mouse microbiome feeding experiment. An open-source implementation of and documentation for SparseDOSSA 2 are available through R/Bioconductor and at http://huttenhower.sph.harvard.edu/sparsedossa2.

## Results

### A statistical model for microbial community profiles

SparseDOSSA is a hierarchical model for microbial count and relative abundance profiles (**Fig. 1**), with components specifically accommodating the major distributional characteristics of such data, namely zero-inflation, compositionality (and thus sequencing depth), feature-feature nonindependence, feature-environment interactions, and high-dimensionality. Briefly (**Fig. 1A**, details in **Methods**), the model a) specifies zero-inflated log-normal marginal distributions for each microbial feature to allow for both biological and technical absences, b) imposes distributions on the “absolute”, i.e. pre-normalized, microbial abundances to satisfy compositionality (similar to models such as the Dirichlet [19] or gamma-MGH [18]), c) models feature-feature correlations through a multivariate Gaussian copula [20], and d) adopts a penalized fitting procedure to address high-dimensionality [21]. Conditional on feature relative abundances and total read depth, count observations are modeled with a standard multinomial sampling procedure, and per-sample read depth is modelled with a log-normal distribution. For implementation, we adopted a penalized Expectation-Maximization procedure for model fitting, and we have evaluated and provided options for cross-validated selection of the optimal penalization parameter (**Methods**). SparseDOSSA 2 is implemented as an R/Bioconductor package (http://huttenhower.sph.harvard.edu/sparsedossa2) and can be accurately fit to a wide variety of different microbial community structures to capture both (inferred) absolute and relative count observations (**Fig. 1B-C**, details below).

**Figure 1:**
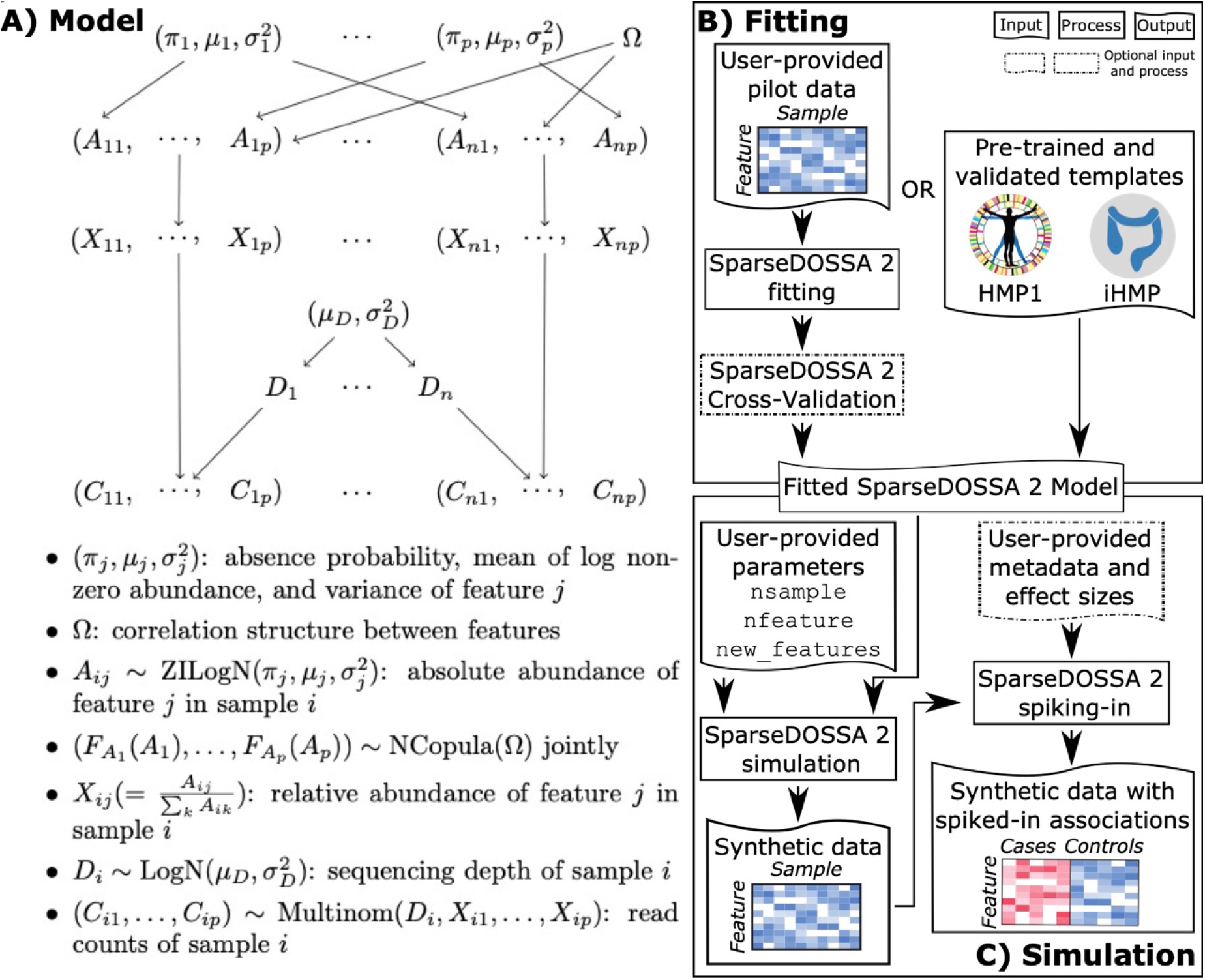
A hierarchical model for microbial community feature profiles. **A)** SparseDOSSA comprises a hierarchical model to capture the generation mechanism of microbial sequencing counts, including components for “hidden” absolute abundances, sequencing depth (and thus compositional relative abundances), zero inflation, and feature-feature and feature-environment interactions. **B)** SparseDOSSA can be fitted to varied microbial community types using cross-validation procedures by users; the software also provides pre-trained models are provided for human microbiome template datasets. This allows for **C)** simulation of either null or “true positive” association spiked-in synthetic datasets, to facilitate microbiome benchmarking or power analysis studies.

### SparseDOSSA accurately recapitulates real-world microbial community structures

We validated SparseDOSSA’s ability to accurately capture realistic microbial community feature profiles by quantifying its performance across a variety of real-world datasets (**Fig. 2**, **Supplemental Table 1**). The studies used include: 1,2) taxonomic profiles from shotgun sequenced metagenomes of healthy human stool and posterior fornix samples from the HMP1-II, hereafter referred to as “Stool” and “Vaginal” [22], 3) shotgun sequenced stool metagenomes of inflammatory bowel disease (IBD) patients from the HMP2 Inflammatory Bowel Disease Multiomics Database (IBDMDB, abbreviated as “IBD”) [2], and 4) 16S rRNA gene sequenced murine distal gut communities after diet perturbation [23]. By evaluating the model in different cohorts, we established its robustness under different community phenotypes, habitats (i.e. body sites), overall ecological structures, and sequencing technologies (**Supplemental Table 1**).

**Figure 2:**
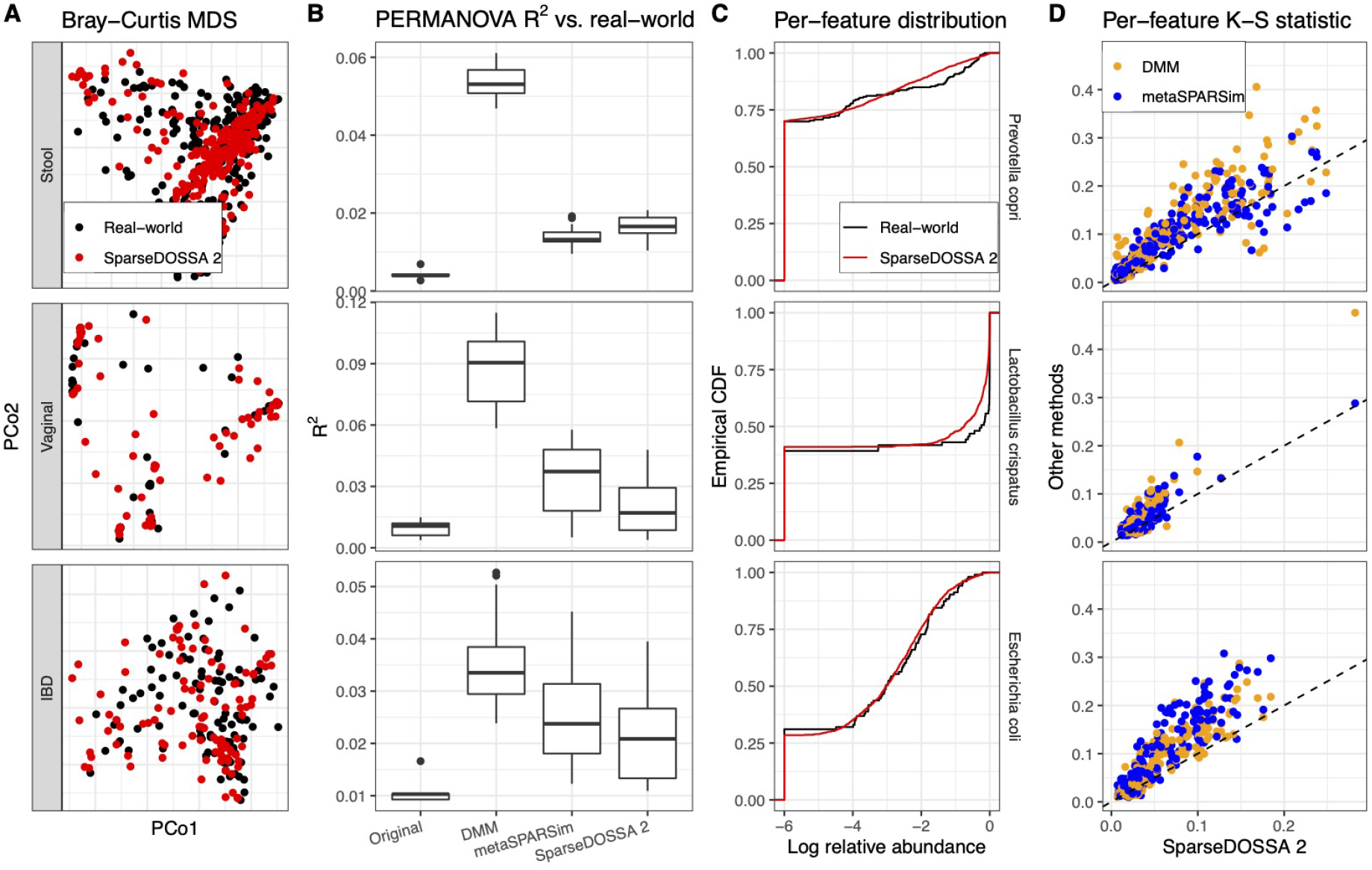
SparseDOSSA accurately recapitulates different microbial community structures. We compared SparseDOSSA 2 simulated microbial counts versus those of three human microbiome training template datasets (Stool, Vaginal, and IBD). **A)** Bray-Curtis ordination shows global agreement between SparseDOSSA 2 simulated microbial abundance profiles and those of their originating real-world populations. **B)** This was quantified by PERMANOVA *R*^2^ statistics, showing that SparseDOSSA 2 simulated samples were significantly less systematically differentiated from their targets than existing DMM and metaSPARSim methods in almost all cases (p-values included in **Supplemental Table 2**). *R*^2^ compared against randomly split original real-world data are included as baseline controls. **C)** Representative features from each environment are similarly distributed between real-world and SparseDOSSA 2 simulated samples, as shown in empirical cumulative distribution functions (CDFs) of log-10 relative abundances (with pseudo value 1e-6 to visually represent zeros). **D)** Per-feature Kolmogorov-Smirnov statistics quantify that SparseDOSSA 2 outperforms existing methods in simulating realistic feature-level relative abundance distributions (p-values are significant and included in **Supplemental Table 3**).

SparseDOSSA 2 captured community parameters and re-simulated microbial profiles with overall community structures that accurately reflected those of the original, real-world ecologies, better than alternative methods (**Fig. 2A-B**), across all human datasets (murine study results reported in separate section). Overall, simulated communities yielded the same patterns of global betadiversity as were contained within each modeled dataset (**Fig. 2A**). This was quantitatively compared against alternative models (Dirichlet-multinomial, DMM [19] and gamma-MGH, namely metaSPARSim [18]) with the PERMANOVA *R*^2^ statistic [24] (**Methods**). We calculated ecological Bray-Curtis dissimilarities between real-world microbial profiles and those simulated by each evaluated method. We then quantified the total variability in the combined dissimilarities that could be attributed to real-world versus simulation difference, expressed as the PERMANOVA *R*^2^. Smaller *R*^2^s thus indicate less deviation of the simulated community structures from the real-world target and better performance of the model.

Across almost all evaluated community types, SparseDOSSA 2 generated significantly smaller *R*^2^ statistics over 25 simulation iterations than existing methods (Mann-Whitney tests p < 0.05), indicating better fit to and recapitulation of the targeted communities (**Fig. 2B,** testing results in **Supplemental Table 2**). Notably, this was consistent in both the human gut (Stool, IBD), where community structure forms continuous “gradients” of microbial composition [9], and the human vaginal environment (Vaginal), where communities are often characterized by a few discrete types with dominant species [25]. Only for the Stool dataset did SparseDOSSA 2 slightly underperform when compared to metaSPARSim in terms of *R*^2^ statistic, while still outperforming with respect to per-feature distributions (**Fig. 2D**). Additionally, metaSPARSim’s simulation procedure can take as much as ~10x longer than SparseDOSSA 2 (**Supplemental Fig. 1**), which is prohibitive for realistic data sizes (especially for benchmarking or power-analysis efforts requiring multiple simulations per parameter configuration, or for Monte-Carlo calculations). We thus conclude that, when evaluated for overall community structures, SparseDOSSA is capable of capturing microbial feature profiles that closely resemble those of real-world microbiomes.

The SparseDOSSA model also provided the best recapitulations of individual features’ relative abundances (**Fig. 2C-D**). For representative features in each environment, the empirical cumulative distribution function (CDF) curves of samples show that SparseDOSSA 2 simulated abundances closely resemble those of the real-world data (**Fig. 2C**). Quantitatively, for each set of microbial features, we measured the difference of distributions between re-simulated and real-world (modeled) relative abundances with the Kolmogorov-Smirnov test statistic (K-S, see **Methods**). The resulting K-S statistic provides a distance between the distribution of each feature’s relative abundances across simulated vs. modeled real-world communities. Smaller K-S statistics thus indicate better performance of the model. SparseDOSSA 2 better approximated the targeted real-world per-feature distributions than existing methods across all evaluated datasets (**Fig. 2D**), reaching statistical significance in each case (**Supplemental Table 3,** Mann-Whitney tests p < 0.05). In addition to simulating existing microbial features, SparseDOSSA 2 also provides the functionality to simulate new features that resemble the targeted environment’s ecological characteristics (**Methods**) and was validated to generate “Stool-like”, “Vaginal-like”, or “IBD-like” new features in terms of prevalence, abundance, and variability for each of the tested datasets (**Supplemental Fig. 2**). Thus both in overall community structure modeling and in per-feature models, SparseDOSSA 2 was able to accurately capture and re-simulate realistic microbial observations better than alternative approaches.

### SparseDOSSA captures covariation among microbes and with real or simulated “phenotypes”

Once the SparseDOSSA model is fit to a real-world microbial community profile, the “reversed” version of the model can be used not only to simulate similar, controlled ecologies, but to spike them with known feature-feature or feature-covariate associations (i.e. metadata “spike-ins”). This is implemented by first capturing the “null” state of targeted real-world studies as described above and by subsequently modifying the fit model parameters to induce artificial associations. Compared to existing spiking-in paradigms [5, 17, 18], the model includes two important improvements (**Methods**). First, SparseDOSSA can model a wide variety of covariates - discrete, continuous, or any combination thereof - with multivariable linear modelling, and can thus accommodate simulations of realistic microbiome population study designs with multiple phenotypes, exposures, or confounders [1]. Second, associations with both non-zero (abundance) and zero-inflated (prevalence) components of microbial features can be captured, along with clearly defined effect sizes (fold change or odds ratio, see **Methods**). This enables rigorous evaluations of, for example, differential abundance testing methods for their statistical performance (e.g. power or false positive rates).

Based on models fit to the Stool and Vaginal communities, SparseDOSSA 2 accurately introduced associations for control “phenotypes” in a new, simulated population (**Fig. 3**, **Supplemental Fig. 3**). Specifically, for the Stool dataset, we introduced a binary covariate (similar to e.g. a case / control contrast) with non-zero effects on 16 (5% of the total 332) microbial features’ abundances (**Fig. 3A**) and prevalences (**Fig. 3B).** Features were selected to ensure the highest effective sample size (**Methods**). For simulated associations of the “phenotype” with feature abundances, log fold change of non-zero relative abundances largely agreed with the target effect sizes within 95% confidence levels (**Fig. 3A**). Prevalence log odds ratios were also as targeted (**Fig. 3B**), with effects in relative abundances mostly agreeing with the prescribed effect sizes. Similar abundance and prevalence results were consistently reproduced in the Vaginal environment (**Supplemental Fig. 3**).

**Figure 3:**
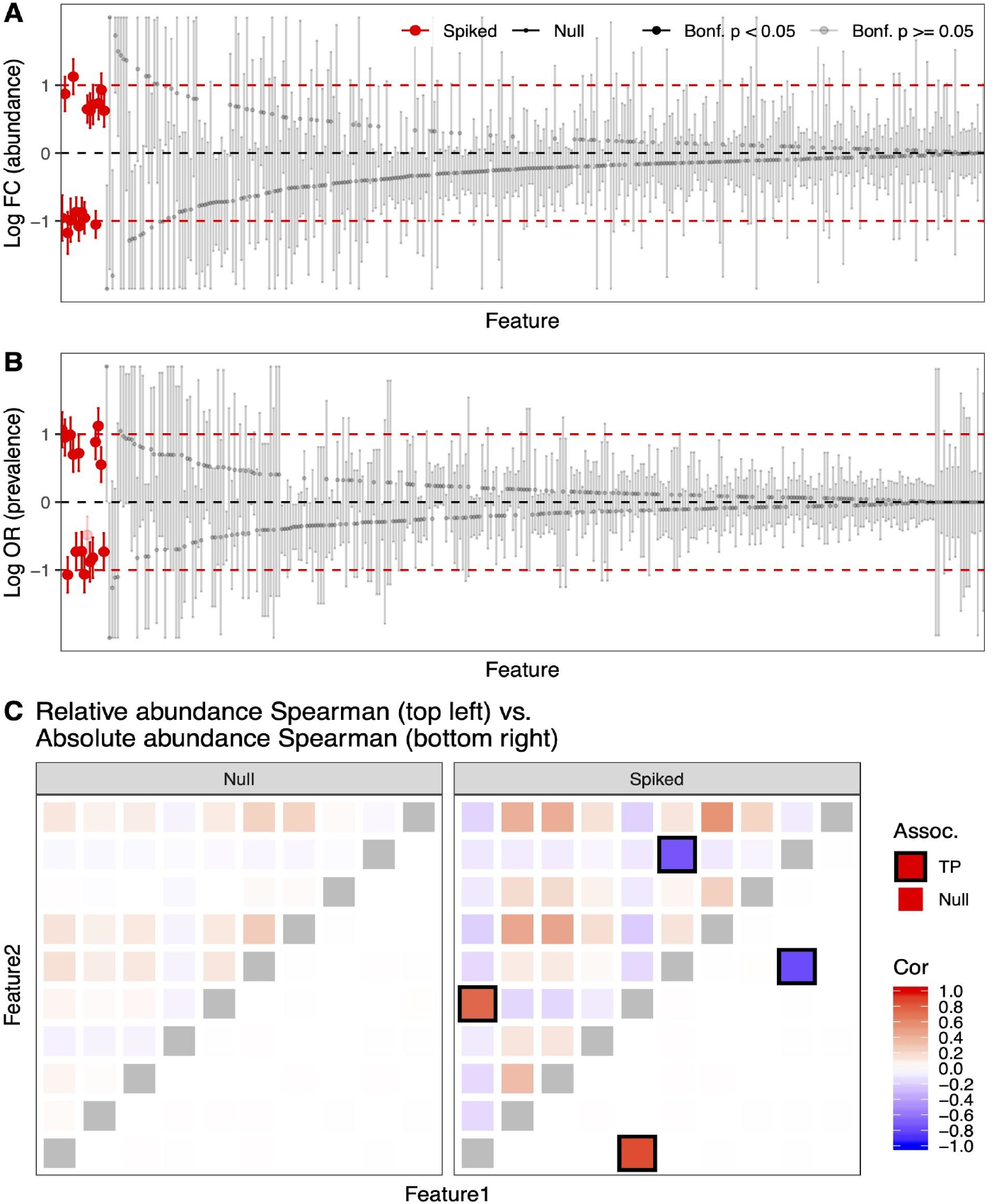
SparseDOSSA can add feature-phenotype and feature-feature associations to modeled microbial community simulations. **A,B** SparseDOSSA 2 correctly simulated feature-phenotype associations targeting the prescribed non-zero relative abundance (**A**) and prevalence (**B**) effect sizes of the spiked features, while maintaining non-associations of null features. **C** SparseDOSSA 2 can also prescribe feature-feature associations, as initially defined in unobserved absolute abundances, which induce correlation structures in simulated relative abundances that correctly include expected spurious correlations caused by compositionality. TP: true positives.

In addition to modeling associations between microbial features and external covariates, SparseDOSSA can also model community ecological interactions (i.e. correlations between microbial features, or feature-feature “spike-ins”, **Fig. 3C**). This is captured by extending the feature-covariate spiking process above to synthetically associate multiple features with the same hidden covariate (**Methods**). First, a null model fit to the Stool community contains no true feature-feature associations, only those that manifest spuriously due to compositionality (**Fig. 3C**). Starting with this, we modified the model to induce increasingly large feature-feature “ecological” interactions (**Supplemental Fig. 4**). SparseDOSSA 2 produced both only and exactly the expected true feature-feature associations among absolute abundance components, and the correct induced compositional correlations after simulating the sequencing assay process (**Fig. 3C**). These results support SparseDOSSA’s ability to modify baseline, null community structures by the introduction of interactions among features or with controlled covariates, which together enable the evaluation of a wide range of statistical approaches to microbiome analysis [15, 26, 27].

### Modeling environment-specific benchmarking and power estimation

Since most microbiome analysis methods make simplifying assumptions that may or may not be suited to particular ecologies, SparseDOSSA’s flexible model enables power and accuracy estimation in a habitat-specific manner (**Fig. 4**). Specifically, by spiking only a limited set of known feature-phenotype associations into an otherwise guaranteed-null model, differential abundance methods can be compared directly to each other in a controlled setting (more details in [28]), enabling targeted method benchmarking (**Fig. 4A**) or power analysis (**Fig. 4B**).

**Figure 4:**
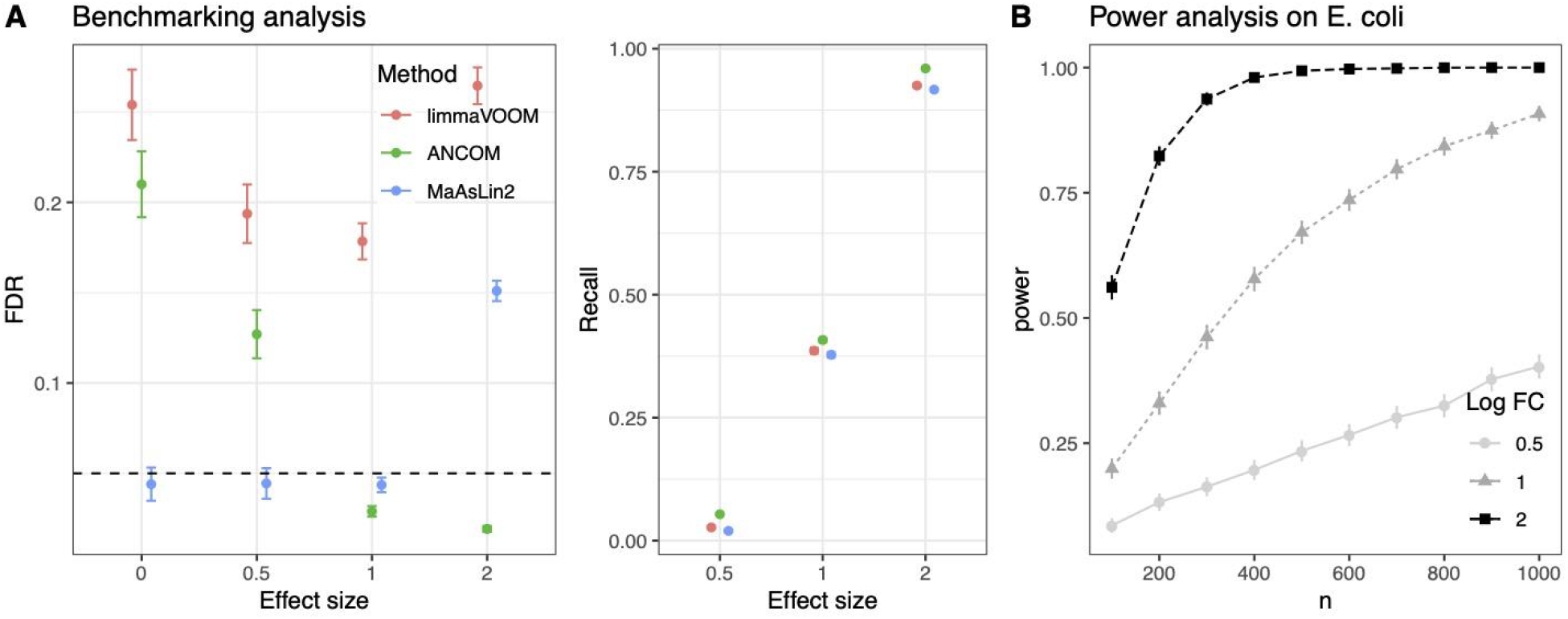
SparseDOSSA enables comparative benchmarking and power analysis of microbial community statistical association tests. For any originating community type of interest, datasets simulated based on a SparseDOSSA model fit can be spiked with known “phenotypes” and feature effect sizes to estimate methods performance (power, FPR, etc.) during (**A**) benchmarking as well as (**B**) power analysis, across controlled combinations of potential effect sizes and sample sizes. Points indicate average performance across simulation repetitions and error bars indicate standard error (**Methods**).

To demonstrate SparseDOSSA’s use for benchmark comparison of microbial community statistical tests, we again simulated synthetic datasets based on the Stool profiles with “phenotypic” associations spiked-in for 5% of features at varying effect sizes (as in **Fig. 3A**). Multiple replicates of the same parameter set were performed to provide performance metric mean and standard errors (**Methods**). Using the resulting gold standards, the performances of three different association tests – limmaVOOM [29], ANCOM [30], and MaAsLin 2 [28] - were similar for power, but false discovery rates varied strikingly (**Fig. 4A**). Notably, the MaAsLin 2 generalized linear model showed good FDR control at small to moderate effect sizes. At higher effect sizes, non spiked-in (“null”) features are also called by MaAsLin 2 as differentially abundant. Interestingly, this is because SparseDOSSA’s spike-in effects are imposed on features’ simulated absolute abundances (**Methods**), and high effect size spike-ins thus also induce relative abundance change in null features due to compositionality. This highlights the important difference between true differential abundance effects corresponding to microbes’ biological variation, versus changes post normalization that are driven by other features.

In contrast, ANCOM [30] was designed to account for compositionality and draw inference about hidden absolute abundances; it successfully and maintained FDR under moderate to strong effect sizes. Arguably as a result, however, its performance suffered for small to null effects, presumably because in such cases it is difficult to distinguish between “driver” microbial features with true absolute effects versus those with changes in their relative abundances due to compositionality. Lastly, limmaVoom [29], designed primarily for RNA-Seq data, had inflated FDRs across all cases.

To demonstrate SparseDOSSA’s use for power analysis during microbial community study design, we focused targeted simulation datasets with spiked-in effects on a feature modeled on *Escherichia coli*, as a microbe commonly associated with dysbiosis in the human gut [2]. Using this approach, SparseDOSSA 2 can be used to estimate each association method’s expected power for similar biomarkers and populations. In this example, MaAsLin 2 has high power to detect a two-fold abundance change in “*E. coli*” for a sample size of at least ~500 individuals, but greatly reduced power for smaller fold-changes (**Fig. 4B**). Since model power for differential abundance testing in sparse, compositional data is extremely difficult to determine parametrically, SparseDOSSA thus provides a way to do so by simulation tailored to any community type or feature of interest.

### SparseDOSSA reproduces an end-to-end diet-microbiome analysis

In many cases, SparseDOSSA thus captures the properties of microbial community ecologies well enough to reproduce surprisingly specific aspects of their membership and distributions, which we next demonstrated by reproducing an end-to-end example from a longitudinal, interventional diet study investigating the effects of diet on the murine gut microbiome [23] (**Fig. 5**). Carmody *et al.* [23] used 16S rRNA gene sequencing to profile changes in the mouse distal gut microbiome under different dietary treatments (chow, raw and cooked tuber, and meat). To determine whether SparseDOSSA could accurately model the microbes, phenotypes, and associations observed over time in these settings, we fitted model parameters for each sample type at different time points and under different treatment assignments (**Methods**).

**Figure 5:**
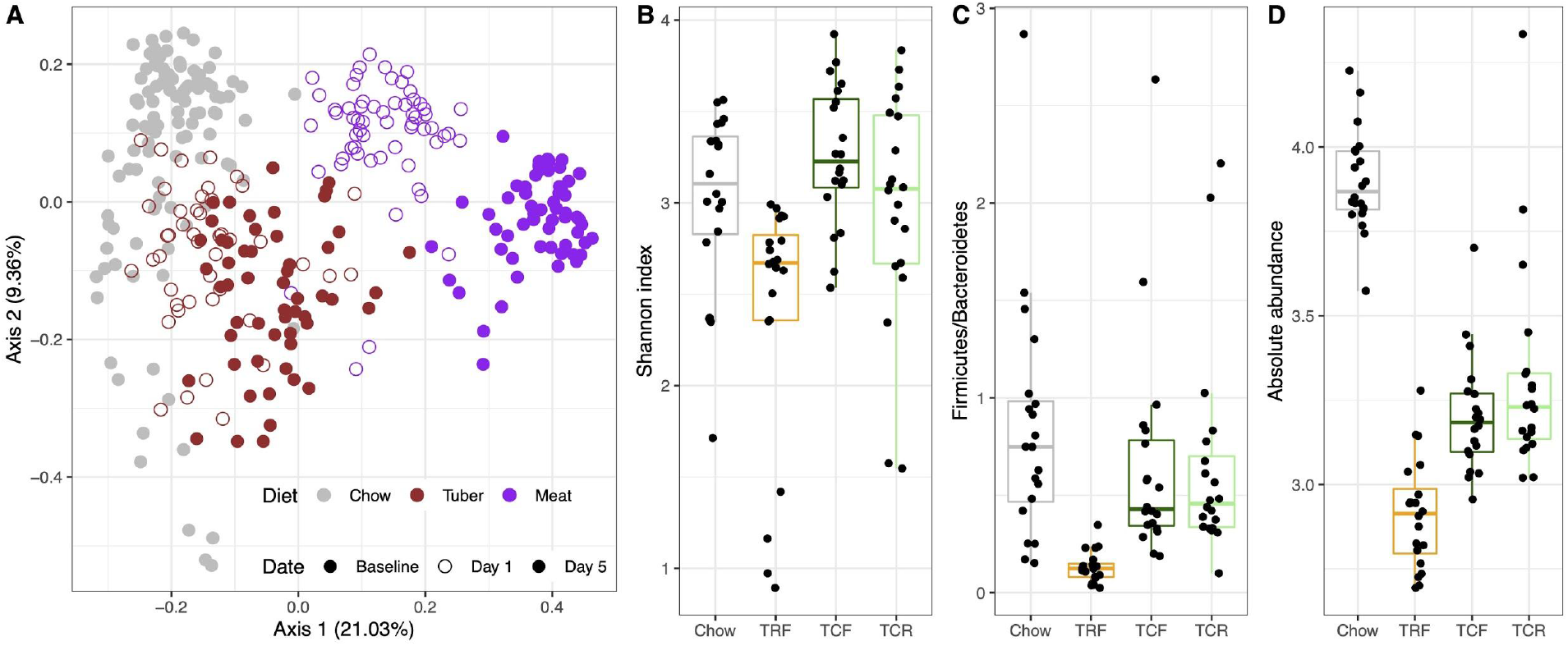
SparseDOSSA correctly models the effects of diet and time on the murine gut microbiome by reproducing effects from amplicon sequencing profiles. **A)** SparseDOSSA 2 was fitted to subsets of samples from [23] that included up to three time points each from collections of mice fed chow, raw or cooked tubers, and meat. The resulting models were then used to simulate controlled microbial community profiles, which correctly reproduced the beta-diversity structures present in the original study (MDS ordination by Bray-Curtis dissimilarities). The SparseDOSSA model was also able to model and synthetically replicate changes in “Bacteroidetes” and “Firmicutes” phyla in response to raw vs. cooked diets, including **B)** overall community alpha-diversity (Shannon index), **C)** the resulting “Firmicutes” vs. “Bacteroidetes” ratio, and **D)** overall whole-community effective biomass. TRF = raw tuber (free-fd); TCF = cooked tuber (free-fed); TCR = cooked tuber (restricted ration).

After re-simulating communities based on these models fits, ordinations of SparseDOSSA 2 results closely mimic the originally observed clustering structure of dietary effects, and even the longitudinal effects of time under treatment (**Fig. 5A**). For quantitative differential abundance effects, based on the observed difference of raw diet (TRF) samples when compared against cooked/free-fed (TCF) and cooked/restricted (TCR) within the tuber diet group [23], we additionally applied SparseDOSSA 2’s spike-in procedure to simulate a ~2x fold increase in the abundance of Bacteroidetes OTUs in TCF/TCR when compared to TCF samples, and a ~2x fold decrease in Firmicutes OTUs (**Methods**). Consequently, the simulated samples displayed similar differential outcomes in community diversity as measured by Shannon index, as well as Firmicutes/Bacteroidetes ratios, as seen in the original study (**Fig. 5B-C**).

Interestingly, even though per-feature absolute abundances are theoretically unidentifiable in the SparseDOSSA model (**Methods**), we note the spiking-procedure recapitulated the decreased total cell counts in TRF (**Fig. 5D**) that [23] also observed via quantitative PCR. The difference between chow versus tuber diets, on the other hand, is completely attributable to SparseDOSSA’s framework, as one can arbitrarily modify the average absolute abundances of our fit to the chow or tuber diet samples, but still yield exactly the same relative abundance profiles. These real-world application results highlight SparseDOSSA’s adaptability to community phenotypes and treatment effects, as well as demonstrate its performance for amplicon sequence datasets and microbial communities associated with non-human hosts.

## Discussion

Here, we have developed a statistical model, implemented in the R package SparseDOSSA 2, for fitting and/or simulating microbial community profiles. These can comprise taxonomic abundances (i.e. relative abundances or counts) from shotgun metagenomic or amplicon sequencing; although not evaluated here, the model is in principle also appropriate for other microbial feature abundances (e.g. genes or pathways). The model can be fit to communities with different host-associated or environmental ecological structures, and it accurately captures their fundamental characteristics, including the distribution of abundances across community members and the diversities of microbial composition across populations. In addition, to support quantitative benchmarking of new methods for microbial community statistics and epidemiology, SparseDOSSA is able to reliably induce user-specified correlation structures involving feature-covariate or feature-feature associations in simulated ecologies. This was demonstrated not only *in silico*, but by end-to-end reproduction of results paralleling those in an interventional mouse feeding study. The underlying generative model thus efficiently and effectively summarizes real microbial communities and recapitulates their latent structure in a manner that is both computationally efficient and statistically principled.

The SparseDOSSA model assumes that the characteristics of a template (real) microbial community are well-captured by the distributions it includes for each component (individual features, feature-feature relationships, sparsity, etc.) More specifically, this requires that 1) the non-zero component of absolute abundances is approximately log-normal, 2) that feature-feature association structure is sparse (as captured by the penalized estimation procedure), and 3) that intrinsic population substructure among samples are absent in the template dataset (i.e. before SparseDOSSA 2 itself optionally spikes-in any such structure). The last assumption 4) that sequencing depths within study are themselves log-normal typically has minimal impact on model fitting or usage. The third assumption holds reasonably well even when any correlation structure originally present is weak or rare relative to overall microbial variance, or affects only a small proportion of features, similar to the assumption of “few differential transcripts” used in most RNA-seq models [31]. Second, inasmuch as the read count of each feature depends on its own observed mean, variance, and sparsity, SparseDOSSA 2’s simulated data will replicate the marginal distribution of the originating template community. This guarantee on the null distribution of subsequently generated communities allows correlation structure (with samples or among features) to be optionally added in isolation for evaluation of microbial community analysis methods. The first assumption is most approximate - it is generally true for ecologically diverse communities, which empirically follow power-law or log-normal behaviors (with a few abundant organisms and a long tail representing the increasingly rare biosphere). However, as discussed above, its violation leads to small residual systematic biases (<0.5%) in communities where tails of rare organisms are more truncated than expected.

Perhaps the greatest strength of the model is its application in simulating microbial community profiles, which we have emphasized and validated here. Most previous methods for associating microbial features with covariates [30, 32–34] or with each other [15, 27, 35, 36] have relied on heterogeneous, one-off models not necessarily reflective of any one “real” microbial community type, or of the diversity of ecological configurations observed in the wild (e.g. the human gut vs. vaginal microbiome vs. soil). By providing a model that can accurately capture many different community types, remove any existing structure through null distributions, and re-introduce known, controlled structure (microbial or covariate), we hope to provide a convenient, unified framework with which statistical methods can be validated specifically for their environments of interest (e.g. human microbiome epidemiology vs. environmental ecological interactions). In addition to this application, while not emphasized here, the model’s parameterization can be used to directly inspect or compare microbial communities. For example, the estimates of absence probabilities *π_j_* for important microbes *j* of interest in specific human populations (e.g. *Prevotella* in the Westernized vs. non-Westernized gut [37]), or the relationships between *π_j_* vs. mean logabundance *μ_j_* across microbes (i.e. prevalence vs. abundance) are directly informative as to their neutral dispersal vs. selection [38]. To some degree this is evident from the murine feeding example above, but most such applications remain to be demonstrated in broader “real-world” datasets.

Relatedly, SparseDOSSA successfully reproduced reported dietary effects on the mouse gut microbiome [23], without assuming such differences *a priori* (**Fig. 5**). By fitting our model on microbial observations of separate treatment groups and time points, we allowed SparseDOSSA to adapt to each subset independently, but without assumptions on the existence or magnitude of differences between them. The emergent reproduction of differentiation by diet in the resulting synthetic communities and features (**Fig. 5A**) exemplifies SparseDOSSA’s utility in capturing environment- or treatment-specific dynamics of real-world microbial communities. In parallel, by introducing effects within each dietary group, SparseDOSSA’s per-feature spike-in procedure was able to reproduce structural microbial community changes such as overall diversity and whole-phylum abundance trade-offs. Together, this end-to-end real-world case study highlights SparseDOSSA’s two key functionalities while also testing a non-human, amplicon-sequenced application context: generating realistic microbial community profiles that closely mimic the targeted environment, and introducing covariate spiked-in microbial perturbations to simulate treatment effects.

With respect to this second use case (covariate effects spike-in), existing simulation models often adopt the simplistic approach of modifying the abundances of taxa in the null community to introduce known associations [5, 18]. SparseDOSSA, in comparison, utilizes rigorous perturbation models to explicitly specify the marginal means of taxa as functions of chosen covariates. This a) enables much more flexible applications such as the inclusion of confounders or random effects (by incorporating them as covariates), and b) yields spiked-in datasets that are strictly compatible with the standard assumptions of (generalized) linear models. Alternatively, differentiation between simple binary (case-versus-control) contrasts could be achieved with our current model by training SparseDOSSA separately on the two corresponding population subsets, given that each was sufficiently large to serve as a template.

Our modeling and simulation procedure for generating feature-feature correlations is, in turn, directly based off the feature-covariate model and comparatively more restrictive; we expect to explore more rigorous and flexible approaches in future work, since any one “correct” way to model ecological associations in absolute vs. relative abundance space is not clear *a priori* [35]. Another related area for future work is in the specific model used for absolute abundances, which are not well-understood from currently available data; our current assumption holds if the total biomass of “typical” communities does not change under “typical” circumstances, but this is obviously quite qualitative. Direct measurements of microbial biomass in some environments such as the human gut have sometimes shown this within approximately one fold change [39, 40], but not in all cases, and certainly not during extreme perturbations such as antibiotics [41].

Thus the SparseDOSSA model simultaneously provides a conceptual framework with which to capture key aspects of microbial ecologies and their members, a simulation system for benchmarking statistical methods that assess correlation structure in microbial community profiles, and a set of marginal parameters for each community and community type of lower dimensionality and potentially reduced noise relative to raw data. The last, while again not yet explored, could allow sample metadata covariates to be more accurately tested for association with microbial features, or tested for association with microbial community features indirectly (e.g. via their prevalence or mean when present). In addition to the areas discussed above, future expansions of the model might include longitudinal structure or other interdependencies among samples (i.e. population substructure), as well as diversifying the application areas for the model (e.g. for power calculations during study design). As currently implemented, SparseDOSSA 2 provides an end-to-end system that enables reproducible and efficient validation of quantitative methods applied to microbial community taxonomic profiles, allowing fair comparisons to be made between different methods or studies to establish a consistent baseline for statistical validation.

## Acknowledgments

This work has been supported in part by National Institutes of Health grants NIDDK R24DK110499 (CH), NIAID U19AI110820 (Owen White), Army Research Office grant W911NF-11-1-0473 (CH), and the Harvard Faculty of Arts and Sciences Dean’s Competitive Fund for Promising Scholarship (LJ).

## Methods

### The SparseDOSSA model

SparseDOSSA uses the following data generation mechanism to parameterize microbial community profiles: a) environments/samples contain microbes with absolute abundances *A*, b) these are normalized to relative abundances *X*, which c) can be measured via sequenced counts *C*. As detailed in **Fig.1A**, our model specification for these components is:

- For the unobserved absolute abundances *A* = (*A*_1_,*A*_2_,⋯,*A_p_*), we specify a Gaussian copula model [20] with zero-inflated log normal marginal distributions. Specifically, this involves assuming hidden multivariate Gaussian variables *g* = (*g*_1_,*g*_2_,⋯,*g_p_*) for the microbial features and a mapping of these variables to the corresponding absolute abundances (*A*_1_,*A*_2_,⋯,*A_p_*):

○ *g* ~ *MVN*(*O, Ω*^−1^). That is, each *g_j_* is a standard *N*(0, 1) variable and their correlation matrix is *Ω*^−1^.
○ Each *g_j_* is mapped to *A_j_* such that *A_j_* follows a zero-inflated log-normal distribution, parameterized by absence probability (*π_j_*) and mean and variability of non-zero log abundances 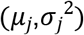:
  ∎ *A_j_* = 0 if *g_j_* < *ϕ*^−1^(*π_j_*)
  ∎ 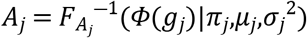 if *g_j_* ≥ *ϕ*^−1^(*π_j_*)
  Where *ϕ* is the standard normal cumulative density function and *F_Aj_* is the cumulative density function of the zero-inflated log-normal distribution, parameterized by 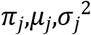. It follows from our model specification that, marginally, *A_j_* follows the prescribed zero-inflated log-normal distribution exactly:

○ With probability *π_j_*, *A_j_* = 0
○ With probability 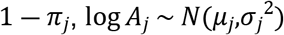 Jointly, correlations between microbial features’ absolute abundances are characterized through the copula parameter *Ω.* The benefit of adopting a copula model is to separate the parameterization and estimation of a joint distribution into its marginal and correlation components; this is illustrated in the **model fitting** subsection below.
- Relative abundances are directly normalized from absolute abundances: 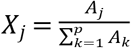. This by definition satisfies compositionality 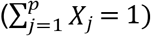. Also note that because *A_j_*’s are zero-inflated, this directly induces zero-inflation (i.e. biological absence) in *X_j_*’s.
- For a given sample *i* with sequencing depth *D_i_*, its per-feature read counts (*C*_1_,*C*_*i*2_,⋯,*C*_*i*p_) are assumed to follow a multinomial distribution with individual features’ probabilities given by *X_ij_*. That is, (*C*_1_,*C*_*i*2_,⋯,*C*_*i*p_) ~ *MultiNom*(*D_i_,X*_*i1*_,*X*_*i*2_,⋯,*X_ip_*), thus also allowing technical zeros.
- Lastly, we assume the sequencing depth *D_i_* across samples follows a log-normal distribution. That is, *D_i_* ~ *LogN*(*μ*_*D*_,*σ*_*D*_^2^).

### Model likelihood

It is helpful to clarify the likelihood of our model given its parameterization. First, we derive *f_A_*, the likelihood for the unobserved absolute abundances *A*. The likelihood of observed data, as we show later, is an integration of *f_A_*. For illustration purposes, we first note the special case where *A_j_* are not zero-inflated. That is, *π_j_* = 0 for all *j*’s. In this case, we have that:

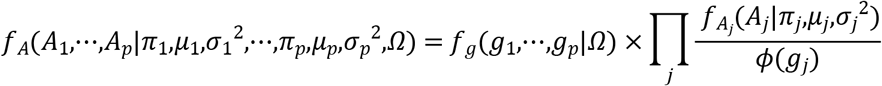

Where *g_j_* is as defined above: *g_j_* = *ϕ*^−1^(*F_A_j__*(*A_j_*)) and *ϕ*(·) is the standard normal density function. The equality follows by noting that the second term (the product) is the Jacobian of the mapping *g*→*A*:*A_j_* = *F_Aj_*^−1^(*ϕ*(*g_j_*)). When one or more *A_j_*’s are zero-inflated, the mapping *g*→*A* is not one-to-one, and the right hand side of the equality requires integration over *g_j_*’s that map to zero-valued *A_j_*’s:

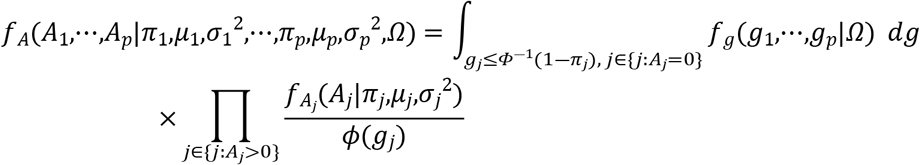

To derive the likelihood for relative abundances *X*, we note that *X*, jointly with the total absolute abundance *A^Σ^* (*A^Σ^*: = *∑_j_ A_j_*), forms a one-to-one mapping with the absolute abundances *A* (*A* = *A^Σ^X*). Thus, the density function for *X, f_x_*, can be obtained through integration of *f_A^∑^,X_*, which is simply *f_A_* multiplied by the Jacobian of the transformation:

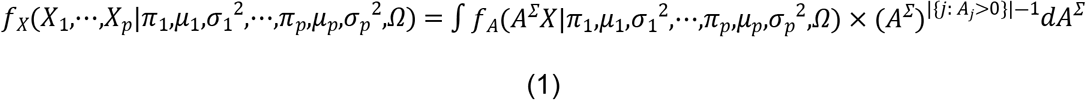

Lastly, for the observed microbial count data *C*, the proper likelihood is:

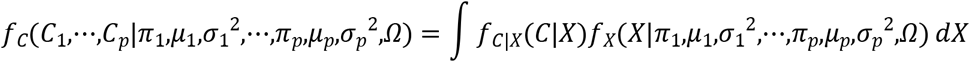

Where *f_C|X_*(*C*) is the multinomial likelihood for microbial counts given their relative abundances. In practice, to simplify computation, during model fitting we replace this likelihood with

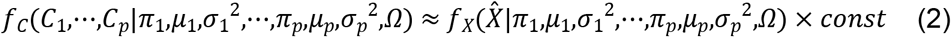

Where 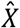 is the multinomial MLE for *X* given observed *C*, i.e., 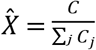, *const* is a normalizing constant not involving the parameters. The approximation is acceptable because with modern sequencing depth [1], *f_C|X_*(*C*|*X*) (as function of *X*) is highly concentrated around 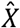. The right-hand side of (2) is what we aim to maximize for estimation of our model’s parameters, 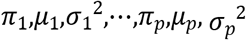, and *Ω*.

### Identifiability

It is important to note that likelihood (1) is unidentifiable. That is, there exist different values of the parameter set (*π*_1_,*μ*_1_,*σ*_1_^2^,⋯,*π_p_,μ_p_,σ_p_*^2^,*Ω*) that yield the same likelihood *f_X_* (and consequently *f_C_*). Intuitively, this is because *X* are normalized from absolute abundances *A*, and different *A* values can map to the same normalized relative abundances *X* - this is thus typical of any compositional setting. Regarding the identifiability of our parameters, we build on the results of [42], which is a special case of our model where *π_j_* = 0 for all *j*’s. Specifically, we note that:

- *π_j_*s are identifiable, as *X_j_* = 0⇔*A_j_*, = 0
- *μ*_1_,⋯,*μ_p_* are identifiable up to a constant. That is, *μ*_1_,⋯,*μ_p_* and *μ*_1_+*c*,⋯,*μ_p_* +*c* lead to the same likelihood, for any constant *c.* For this reason, in our model estimation we impose the (arbitrary) constraint that *∑_j_ μ_j_* = 0.
- *σ*_1_^2^,⋯,*σ_p_*^2^ are identifiable, given *μ*_1_,⋯,*μ_p_* and *Ω*. One can note that when *π_j_* = 1 for all *j*’s, our likelihood degenerates to that in [42] with explicit analytical forms.
- *Ω* is not identifiable. Again, consider the special case that *π_j_* = 1 and *σ_j_* = 1, the form of *f_X_* is explicit and involves 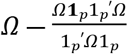, which is a multiple-to-one mapping from *Ω.* The issue of non-identifiable correlation matrices for microbiome abundance data has been noted and addressed in many previous works; refer to {25950956; 28489411; 29140991; doi.org/10.1080/01621459.2018.1442340} for a partial list. We adopt the technique used in many of these previous works, namely *L*_1_ penalization on *Ω*, to simultaneously address the identifiability issue as well as high-dimensionality for generic estimation of large covariance matrices [21].

### Model fitting

Given our model specification and its (non-)identifiability, we propose to minimize the following penalized negative log-likelihood function for solving the parameter set 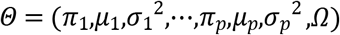 (the sequencing depth parameters (*μ_D_,σ_D_*^2^) can be fitted independently on per-sample read depths with maximum likelihood):

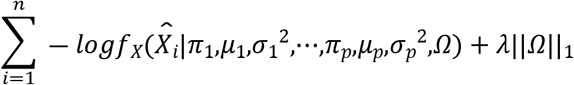

Subject to the constraint for *μ_j_* as specified above: *∑_j_ μ_j_* ⋯ = 0. As such, *λ* >0 is a penalizing tuning parameter, which we choose with cross-validation in practice. 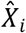 can be either existing relative abundance estimations or, as specified above, normalized from count observations 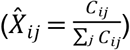.

As specified in (1), the likelihood function *f_X_* involves integration over *A^Σ^* and is not analytically tractable. Numerically, we propose the following penalized expectation-maximization algorithm [43] for model fitting:

1. Initialize 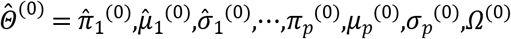 by fitting a multivariate log-normal distribution on 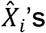.
2. During the *r*-th iteration:

a. E-step: calculate expectation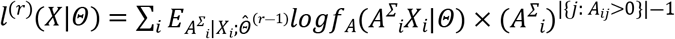
b. Penalized M-step: maximize 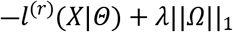 to obtain 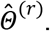 Note that 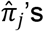 do not require updates. 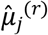 and 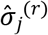 can be solved analytically. 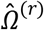 can be solved with standard graphical lasso [21].
3. Iterate until convergence.

### Generating synthetic microbial observations and simulating new features

Given that our model is fully parametric, synthetic microbial observations, including (hidden) absolute abundances, normalized relative abundances, and sequencing counts, can be generated following the same specifications as described above. To provide model parameters, the user can adopt one of the pre-trained sets included with the software or use the SparseDOSSA 2 training procedure to estimate parameters from any microbial template dataset suited for their simulation case.

Users may also be interested in generating “new” microbial features from the same ecological environment. For this, SparseDOSSA additionally models the per-feature parameters (*π_j_,μ_j_,σ_j_*^2^) with a three-dimensional non-parametric distribution *F.* That is, across features, (*π_j_,μ_j_,σ_j_*^2^) ~ *F.* Given a set of SparseDOSSA fitted results 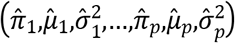, *F* can be estimated with a threedimensional normal kernel density estimator [44]. The estimated 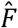 can then be used to simulate new microbial features that follow the ecological characteristics (i.e., prevalence, abundance, and variability) of the fitted environment (**Supplemental Fig. 2**).

### Association spike-in

SparseDOSSA adopts linear and generalized linear models for flexible spiking-in in both microbial features’ non-zero abundances and prevalences, based on covariates. Let *Z_i_* be the vector of covariate(s) for sample *ι* and *β* be the targeted corresponding effect sizes (coefficients). To spike in associations between feature *j*’s abundance and covariates *Z_i_*, we modify the feature’s nonzero mean log absolute abundance parameter *μ_j_* across samples. Specifically, the post spike-in mean log abundance is modified as

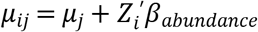

For the *i*-th sample, *A_ij_* can be generated with 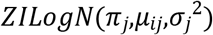 instead of the original 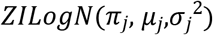. This dictates that *A_ij_*’s are associated with *Z_i_* in their mean non-zero log abundances. As *μ_ij_* are specified on the log scale, *β_abundance_*, by definition, corresponds to log fold changes.

The prevalence spike-in similarly is specified via the logistic model; we modify the presence probability parameter (1 — *π_j_*) across samples:

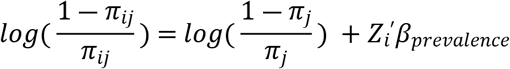

And generate *A_ij_*’s correspondingly. This introduces an association between the covariates *Z_i_* and feature *j*’s prevalence, with *β_prevalence_* corresponding to log odds ratios of the feature being present. The multivariate linear modelling approach for specifying the association effects for both abundance and prevalence allows us to flexibly incorporate different variable types (e.g. binary, continuous, etc.) and study designs (e.g. existence of confounders).

We note that, importantly, our spiking-in procedure is performed on the absolute abundances, A, which induces differential effects in relative abundances *X* (**Fig. 3A-B**). The main benefit of this approach is that both the spiked-in microbial features and the “null” (i.e. non-spiked features) are clearly defined. The alternative - specifying effects for *X_j_* - is conceptually difficult. As *X* is compositional (sums to 1), prescribing enrichment effects (higher abundance or prevalence) for some microbial features must by definition lead to depletion effects for certain other features. This renders it difficult to clearly define and separate the set of “true positive” spiked-in microbial features and the set of null features. SparseDOSSA’s definition of effects for absolute abundances in its spike-in procedure align with recent efforts to rigorously characterize microbial differential abundance effects under the constraint of compositionality [30, 45]. Empirically, we note that prescribed log fold changes or odds ratios for *Aj* often lead to similar effect sizes in the relative abundances *X_j_* for the spiked-in feature *j*’s (**Fig. 3A-B**).

Lastly, we note that the spiking-in procedure with metadata variables can be used to simulate association effects between pairs of microbial features (**Fig. 3C**). Specifically, we first simulate a hidden covariate *Z* with standard normal distribution. For a pair of features *j*_1_, and *j*_2_, to enforce positive correlations between the two absolute abundances *A*_*j*1_ and *A*_*j*2_, we simulate for them to be associated with *Z* in the same direction:

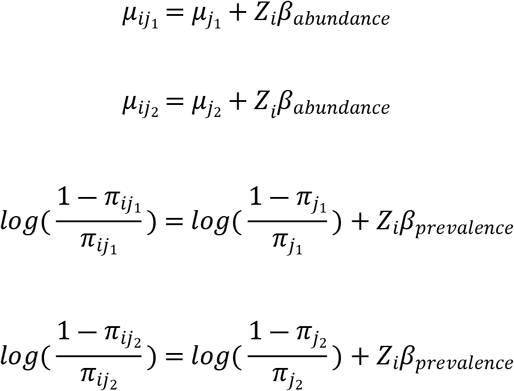

Where *β_abundance_* = *β_prevalence_* = *β* can be viewed here as the effect size specifying the strength of correlation between *A*_*j*1_ and *A*_*j*2_ To spike in negative correlation between the two, we simply keep *β* as the effect for one of the features and use −*β* for the other.

### Evaluation with real-world datasets

For evaluation and comparison of microbiome simulation methods, we examined three real-world datasets with different host environments and disease statuses (**Supplemental Table 1**) [2, 22]. We subset publicly available species level profiles from [22] (all healthy) to baseline time point stool (Stool) and posterior fornix (Vaginal) samples, and those from [2] to baseline time point IBD samples (IBD, including Crohn’s disease and ulcerative colitis patients). We additionally removed samples with lower than ~3,000 reads mapped to identified taxa and features present in less than 3 samples. The datasets’ dimensions (sample size, number of features), post processing and filtering, are included in **Supplemental Table 1**.

To evaluate the performance of individual methods, we randomly partitioned each dataset (Stool, Vaginal, IBD) into halves for five iterations. For each partitioning, we fit the parameterization/simulation methods (DMM, metaSPARSim, and SparseDOSSA 2) on one half of the data (training). We then simulated synthetic microbial observations with the same sample size using the fitted results. Lastly, we compared the synthetic observations with the other half of the partitioned data (testing) in terms of both overall dissimilarity with PERMANOVA [24] and per-feature distribution differences (methods detailed below). Within each partitioning, this simulation was also performed five times for each method. That is, each method was used to randomly simulate five synthetic datasets for comparison with the testing half. The partitioning procedure allows us to evaluate method performance without a model overfitting effect. The DMM was fitted using R package “dirmult” and metaSPARSim was fitted using the implementation referred to in its publication [18]. For metaSPARSim fitting, the percent not zero filter for features was set to 0 instead of the default 0.2. In our evaluation this led to an observed performance increase (thus a favorable assessment), likely due to the existence of many highly zero inflated microbial species.

To evaluate overall dissimilarity between the original and synthetic samples, for each partitioning we combined the testing half of the original samples with the simulated datasets (five for each fitted method). We calculated the sample-to-sample Bray-Curtis dissimilarity matrix *D* on the combined dataset. The univariate PERMANOVA model, *D* ~ *I*{*Sample is simulated*} was fitted. The corresponding *R*^2^ statistic quantifies the percentage of variability between samples attributable to the difference between original “real-world” as compared to simulated samples. Smaller *R*^2^ statistics indicate less difference, and better performance of the simulation method. For each method, a total of 25 PERMANOVA evaluations (5 original dataset partitioning × 5 simulation) was performed for each real-world dataset. Lastly, we additionally evaluated the *R*^2^ between the training and testing halves of a dataset for each partitioning; this yields an estimation of minimum achievable β^2^’s for each dataset.

To evaluate the difference between distributions of individual features in the original and synthetic datasets, we simply combined the synthetic datasets generated across all partitioning and simulation repetitions. An individual feature was compared for its relative abundance distribution between the original real-world data and combined synthetic samples. This was quantified with the Kolmogorov-Smirnov (K-S) test statistic, which is defined as the largest absolute difference between the empirical cumulative distribution functions of the real-world and synthetic abundances. Smaller K-S statistics indicate better approximation of the targeted real-world distributions with the simulation method.

### Association spike-in evaluation

We simulated spiked-in associations between microbial features and a synthetic case/control variable, based on the SparseDOSSA 2 fitted results. A total of 1,000 synthetic samples were simulated (500 cases and 500 controls). For non-zero abundance spike-in (**Fig. 3A**), the top 5% (16 total) most prevalent features were selected for spiking-in; this yields the highest effective sample size for the selected features because our abundance spiking-in targets only the non-zero component of a feature’s distribution. Half of the features were spiked for a targeted log fold change (*β_abundance_*) of 1 in cases compared to controls, and the other half were spiked for a log fold change of −1. Actual log fold changes in the simulated relative abundances, along with 95% confidence intervals, were calculated by performing a linear regression on the log transformed non-zero relative abundances for each feature.

Similarly, for prevalence spiking-in (**Fig. 3B**), the top 5% features with prevalence closest to 0.5 were selected; as with abundance spike-ins, this was to ensure the spiked-in features had the highest effective sample size, as the association between a binary outcome (presence/absence here) and a binary covariate (case/control) is best-powered when the sample distribution is balanced across all different outcome/covariate combinations [46]. Again, half of these features were spiked for a targeted log odds ratio (*β_prevalence_*) of 1 in cases compared to controls, and the other half were spiked for a log odds ratio of −1. Actual log odds ratios of the simulated feature prevalence, along with 95% confidence intervals, were calculated by performing a logistic regression on the presence for each feature.

For simulation of feature-feature associations, we first set the correlation between feature pairs in SparseDOSSA 2 to zero (i.e., *Ω* = I where *I* is the identity matrix). This ensures that feature absolute abundances are independent in the “null” dataset (**Fig. 3C** left panel, bottom right), whereas spurious correlation still exists in relative abundances due to compositionality. Two random pairs (four features) in the top ten most abundant features were selected for non-zero feature-feature association spike-in. As specified in **Methods** above, we simulated two independent normal synthetic hidden metadata variables, one for each feature pair to be associated. For the first feature pair, they were spiked in both abundance and prevalence with the same effect (*β* = *β_abundance_* = *β_prevalence_*) at varying sizes, for positive association. The second pair were spiked with opposing effects (*β* for one, −*β* for the other) for negative association. We used Spearman correlation to estimate the empirical association between feature pairs in the simulated absolute and relative abundances. Target association effect size was also varied (*β* of 0, 1, 2, and 5) to showcase the relative signals of “true” associations that exist for both absolute and relative abundances, and spurious associations that are only induced in relative abundances due to compositionality (**Fig. 3C**, **Supplemental Fig. 4**)

### Benchmarking and power analysis

Since “true” associations with prescribed effect sizes are known for SparseDOSSA synthetic datasets, they can be used for benchmarking microbiome analysis methods as well as for power analysis of microbiome study designs. For benchmarking analysis (**Fig. 4A**), we again selected the top 5% (16 total) most prevalent features in the Stool dataset to perform abundance spike-in, such that the selected features had the highest effective sample size. A total of 200 microbial profiles were simulated to be associated with a balanced binary metadata (100 cases, 100 controls). We varied effect sizes with half spiked features at *β_abundance_* = (0, 0.5, 1, 2) and the other half with *β_abundance_* = (0, – 0.5, – 1, – 2), correspondingly (in the effect size 0 case no spike-in was performed and microbial profiles are generated independently of metadata). A total of 500 random simulations were performed for each parameter combination. We applied existing differential abundance analysis methods to detect the spiked-in features in each simulation dataset [28–30], with individual method configurations as reported in our previous benchmarking analysis [28]. We summarized the empirical power and FDR of a method in one simulation dataset, across the twenty random replicates for each parameter configuration, and reported the mean and standard error in **Fig. 4A**.

For showcasing SparseDOSSA’s utility in a power analysis, we spiked in non-zero abundance associations with a balanced case-control variable for a simulated species parameterized by fitting *Escherichia coli.* This was performed at varying effect sizes (log fold change, *β_abundance_* = (0.5, 1, 2)) and sample sizes (100 to 1000). For each parameter configuration, a total of 500 replicates were simulated. The empirical power and its standard error of using MaAsLin 2 to detect the differential abundant effect in “*E. coli*” was summarized across the 500 replicates and reported in **Fig. 4B**; this was repeated for each effect size/sample size configuration.

### Murine diet microbiome analysis

We applied SparseDOSSA 2 to the longitudinal diet dataset of the mouse gut microbiome in [23], to show that our method is capable of reproducing a complex study’s findings. To recapitulate the longitudinal diet effect as reported in [23]’s Fig. 1a, we fitted SparseDOSSA 2 separately on 1) the control Chow diet samples at baseline, 2,3) Tuber diet samples at day 1 and day 5, separately, and 4,5) Meat diet samples at day 1 and day 5, separately. This approach allows SparseDOSSA 2 to independently fit subsets of the data, without assuming *a priori* the observed differences noted in [23]. We then used SparseDOSSA 2 fitted results to simulate synthetic observations for each diet/timepoint combination, with five times the original sample size (to reduce variability due to random sampling). Bray-Curtis MDS ordination on these synthetic data displayed a striking resemblance to that observed in [23] (**Fig. 5A**), in that a) communities cluster according to dietary treatment, and 2) this response is consistent after one day of switching from chow to whole-food diets and is strengthened at day 5.

We next reproduced the differential gut microbial profiles observed in mice fed raw versus tuber diets as presented in [23]’s Fig. 1f-g. [23] adopted three different types of Tuber diet: the raw/free-fed (TRF), the cooked/free-fed (TCF), and the cooked/restricted (TCR). This study presented that on the phylum level, TRF induced enrichment of Bacteroidetes and depletion of Firmicutes when compared to TCF/TCR. We applied SparseDOSSA 2’s feature spike-in procedure to approximate this effect. Specifically, we generated a balanced, three category (TRF/TCF/TCR) variable. Based on our fitted model of the Tuber diet at day 5, we spiked in a two-fold (*β_abundance_* = log 2) increase in the non-zero abundance of Bacteroidetes OTUs and a two-fold decrease in Firmicutes OTUs (*β_abundance_* = – log 2) in TRF samples when compared to TCF/TCR samples. This roughly agrees with the presented results in [23] Fig. 1d. We next simulated SparseDOSSA synthetic datasets for both the baseline Chow diet samples (sample size = 20), and the spiked-in Tuber diet samples (60 samples total, 20 each for TRF/TCF/TCR). We calculated the Shannon index and Firmicutes/Bacteroidetes ratios of these samples, and show in **Fig. 5B** that they agreed with the corresponding findings presented in [23] Fig. 1f-g.

## References

1. Mallick H, Ma S, Franzosa EA, Vatanen T, Morgan XC, Huttenhower C. Experimental design and quantitative analysis of microbial community multiomics. Genome Biol. 2017;18(1):228. Epub 2017/12/01. doi: 10.1186/s13059-017-1359-z. PubMed PMID: 29187204; PubMed Central PMCID: PMCPMC5708111.

2. Lloyd-Price J, Arze C, Ananthakrishnan AN, Schirmer M, Avila-Pacheco J, Poon TW, et al. Multi-omics of the gut microbial ecosystem in inflammatory bowel diseases. Nature. 2019;569(7758):655–62. Epub 2019/05/31. doi: 10.1038/s41586-019-1237-9. PubMed PMID: 31142855; PubMed Central PMCID: PMCPMC6650278.

3. Wirbel J, Pyl PT, Kartal E, Zych K, Kashani A, Milanese A, et al. Meta-analysis of fecal metagenomes reveals global microbial signatures that are specific for colorectal cancer. Nat Med. 2019;25(4):679–89. Epub 2019/04/03. doi: 10.1038/s41591-019-0406-6. PubMed PMID: 30936547.

4. Gloor GB, Macklaim JM, Pawlowsky-Glahn V, Egozcue JJ. Microbiome Datasets Are Compositional: And This Is Not Optional. Front Microbiol. 2017;8:2224. Epub 2017/12/01. doi: 10.3389/fmicb.2017.02224. PubMed PMID: 29187837; PubMed Central PMCID: PMCPMC5695134.

5. McMurdie PJ, Holmes S. Waste not, want not: why rarefying microbiome data is inadmissible. PLoS Comput Biol. 2014;10(4):e1003531. Epub 2014/04/05. doi: 10.1371/journal.pcbi.1003531. PubMed PMID: 24699258; PubMed Central PMCID: PMCPMC3974642.

6. Qin J, Li Y, Cai Z, Li S, Zhu J, Zhang F, et al. A metagenome-wide association study of gut microbiota in type 2 diabetes. Nature. 2012;490(7418):55–60. Epub 2012/10/02. doi: 10.1038/nature11450. PubMed PMID: 23023125.

7. Forslund K, Hildebrand F, Nielsen T, Falony G, Le Chatelier E, Sunagawa S, et al. Disentangling type 2 diabetes and metformin treatment signatures in the human gut microbiota. Nature. 2015;528(7581):262–6. Epub 2015/12/04. doi: 10.1038/nature15766. PubMed PMID: 26633628; PubMed Central PMCID: PMCPMC4681099.

8. Sinha R, Abu-Ali G, Vogtmann E, Fodor AA, Ren B, Amir A, et al. Assessment of variation in microbial community amplicon sequencing by the Microbiome Quality Control (MBQC) project consortium. Nat Biotechnol. 2017;35(11):1077–86. Epub 2017/10/03. doi: 10.1038/nbt.3981. PubMed PMID: 28967885; PubMed Central PMCID: PMCPMC5839636.

9. Koren O, Knights D, Gonzalez A, Waldron L, Segata N, Knight R, et al. A guide to enterotypes across the human body: meta-analysis of microbial community structures in human microbiome datasets. PLoS Comput Biol. 2013;9(1):e1002863. Epub 2013/01/18. doi: 10.1371/journal.pcbi.1002863. PubMed PMID: 23326225; PubMed Central PMCID: PMCPMC3542080.

10. Tusher VG, Tibshirani R, Chu G. Significance analysis of microarrays applied to the ionizing radiation response. Proc Natl Acad Sci U S A. 2001;98(9):5116–21. Epub 2001/04/20. doi: 10.1073/pnas.091062498. PubMed PMID: 11309499; PubMed Central PMCID: PMCPMC33173.

11. Irizarry RA, Hobbs B, Collin F, Beazer-Barclay YD, Antonellis KJ, Scherf U, et al. Exploration, normalization, and summaries of high density oligonucleotide array probe level data. Biostatistics. 2003;4(2):249–64. Epub 2003/08/20. doi: 10.1093/biostatistics/4.2.249. PubMed PMID: 12925520.

12. Nykter M, Aho T, Ahdesmaki M, Ruusuvuori P, Lehmussola A, Yli-Harja O. Simulation of microarray data with realistic characteristics. BMC Bioinformatics. 2006;7:349. Epub 2006/07/20. doi: 10.1186/1471-2105-7-349. PubMed PMID: 16848902; PubMed Central PMCID: PMCPMC1574357.

13. Park T, Yi SG, Kang SH, Lee S, Lee YS, Simon R. Evaluation of normalization methods for microarray data. BMC Bioinformatics. 2003;4:33. Epub 2003/09/03. doi: 10.1186/1471-21054-33. PubMed PMID: 12950995; PubMed Central PMCID: PMCPMC200968.

14. Huang W, Li L, Myers JR, Marth GT. ART: a next-generation sequencing read simulator. Bioinformatics. 2012;28(4):593–4. Epub 2011/12/27. doi: 10.1093/bioinformatics/btr708. PubMed PMID: 22199392; PubMed Central PMCID: PMCPMC3278762.

15. Schwager E, Mallick H, Ventz S, Huttenhower C. A Bayesian method for detecting pairwise associations in compositional data. PLoS Comput Biol. 2017;13(11):e1005852. Epub 2017/11/16. doi: 10.1371/journal.pcbi.1005852. PubMed PMID: 29140991; PubMed Central PMCID: PMCPMC5706738.

16. Paulson JN, Stine OC, Bravo HC, Pop M. Differential abundance analysis for microbial marker-gene surveys. Nat Methods. 2013;10(12):1200–2. Epub 2013/10/01. doi: 10.1038/nmeth.2658. PubMed PMID: 24076764; PubMed Central PMCID: PMCPMC4010126.

17. Thorsen J, Brejnrod A, Mortensen M, Rasmussen MA, Stokholm J, Al-Soud WA, et al. Large-scale benchmarking reveals false discoveries and count transformation sensitivity in 16S rRNA gene amplicon data analysis methods used in microbiome studies. Microbiome. 2016;4(1):62. Epub 2016/11/26. doi: 10.1186/s40168-016-0208-8. PubMed PMID: 27884206; PubMed Central PMCID: PMCPMC5123278.

18. Patuzzi I, Baruzzo G, Losasso C, Ricci A, Di Camillo B. metaSPARSim: a 16S rRNA gene sequencing count data simulator. BMC Bioinformatics. 2019;20(Suppl 9):416. Epub 2019/11/24. doi: 10.1186/s12859-019-2882-6. PubMed PMID: 31757204; PubMed Central PMCID: PMCPMC6873395.

19. Chen J, Li H. Variable Selection for Sparse Dirichlet-Multinomial Regression with an Application to Microbiome Data Analysis. Ann Appl Stat. 2013;7(1). Epub 2013/12/07. doi: 10.1214/12-AOAS592. PubMed PMID: 24312162; PubMed Central PMCID: PMCPMC3846354.

20. Murray JS, Dunson DB, Carin L, Lucas JE. Bayesian Gaussian Copula Factor Models for Mixed Data. J Am Stat Assoc. 2013;108(502):656–65. Epub 2013/08/31. doi: 10.1080/01621459.2012.762328. PubMed PMID: 23990691; PubMed Central PMCID: PMCPMC3753118.

21. Friedman J, Hastie T, Tibshirani R. Sparse inverse covariance estimation with the graphical lasso. Biostatistics. 2008;9(3):432–41. Epub 2007/12/15. doi: 10.1093/biostatistics/kxm045. PubMed PMID: 18079126; PubMed Central PMCID: PMCPMC3019769.

22. Lloyd-Price J, Mahurkar A, Rahnavard G, Crabtree J, Orvis J, Hall AB, et al. Strains, functions and dynamics in the expanded Human Microbiome Project. Nature. 2017;550(7674):61–6. Epub 2017/09/28. doi: 10.1038/nature23889. PubMed PMID: 28953883; PubMed Central PMCID: PMCPMC5831082.

23. Carmody RN, Bisanz JE, Bowen BP, Maurice CF, Lyalina S, Louie KB, et al. Cooking shapes the structure and function of the gut microbiome. Nat Microbiol. 2019;4(12):2052–63. Epub 2019/10/02. doi: 10.1038/s41564-019-0569-4. PubMed PMID: 31570867; PubMed Central PMCID: PMCPMC6886678.

24. Tang ZZ, Chen G, Alekseyenko AV. PERMANOVA-S: association test for microbial community composition that accommodates confounders and multiple distances. Bioinformatics. 2016;32(17):2618–25. Epub 2016/05/21. doi: 10.1093/bioinformatics/btw311. PubMed PMID: 27197815; PubMed Central PMCID: PMCPMC5013911.

25. Ravel J, Gajer P, Abdo Z, Schneider GM, Koenig SS, McCulle SL, et al. Vaginal microbiome of reproductive-age women. Proc Natl Acad Sci U S A. 2011;108 Suppl 1:4680–7. Epub 2010/06/11. doi: 10.1073/pnas.1002611107. PubMed PMID: 20534435; PubMed Central PMCID: PMCPMC3063603.

26. Cao Y, Lin W, Li H. Large covariance estimation for compositional data via composition-adjusted thresholding. Journal of the American Statistical Association. 2019;114(526):759–72. doi: doi.org/10.1080/01621459.2018.1442340.

27. Kurtz ZD, Muller CL, Miraldi ER, Littman DR, Blaser MJ, Bonneau RA. Sparse and compositionally robust inference of microbial ecological networks. PLoS Comput Biol. 2015;11(5):e1004226. Epub 2015/05/08. doi: 10.1371/journal.pcbi.1004226. PubMed PMID: 25950956; PubMed Central PMCID: PMCPMC4423992.

28. Mallick H, Rahnavard A, McIver LJ, Ma S, Zhang Y, Nguyen LH, et al. Multivariable Association Discovery in Population-scale Meta-omics Studies. bioRxiv. 2021:2021.01.20.427420. doi: 10.1101/2021.01.20.427420.

29. Law CW, Chen Y, Shi W, Smyth GK. voom: Precision weights unlock linear model analysis tools for RNA-seq read counts. Genome Biol. 2014;15(2):R29. Epub 2014/02/04. doi: 10.1186/gb-2014-15-2-r29. PubMed PMID: 24485249; PubMed Central PMCID: PMCPMC4053721.

30. Mandal S, Van Treuren W, White RA, Eggesbo M, Knight R, Peddada SD. Analysis of composition of microbiomes: a novel method for studying microbial composition. Microb Ecol Health Dis. 2015;26:27663. Epub 2015/06/02. doi: 10.3402/mehd.v26.27663. PubMed PMID: 26028277; PubMed Central PMCID: PMCPMC4450248.

31. Zhou YH, Xia K, Wright FA. A powerful and flexible approach to the analysis of RNA sequence count data. Bioinformatics. 2011;27(19):2672–8. Epub 2011/08/04. doi: 10.1093/bioinformatics/btr449. PubMed PMID: 21810900; PubMed Central PMCID: PMCPMC3179656.

32. Morgan XC, Tickle TL, Sokol H, Gevers D, Devaney KL, Ward DV, et al. Dysfunction of the intestinal microbiome in inflammatory bowel disease and treatment. Genome Biol. 2012;13(9):R79. Epub 2012/09/28. doi: 10.1186/gb-2012-13-9-r79. PubMed PMID: 23013615; PubMed Central PMCID: PMCPMC3506950.

33. Zhao N, Chen J, Carroll IM, Ringel-Kulka T, Epstein MP, Zhou H, et al. Testing in Microbiome-Profiling Studies with MiRKAT, the Microbiome Regression-Based Kernel Association Test. Am J Hum Genet. 2015;96(5):797–807. Epub 2015/05/11. doi: 10.1016/j.ajhg.2015.04.003. PubMed PMID: 25957468; PubMed Central PMCID: PMCPMC4570290.

34. McMurdie PJ, Holmes S. phyloseq: an R package for reproducible interactive analysis and graphics of microbiome census data. PLoS One. 2013;8(4):e61217. Epub 2013/05/01. doi: 10.1371/journal.pone.0061217. PubMed PMID: 23630581; PubMed Central PMCID: PMCPMC3632530.

35. Weiss S, Van Treuren W, Lozupone C, Faust K, Friedman J, Deng Y, et al. Correlation detection strategies in microbial data sets vary widely in sensitivity and precision. ISME J. 2016;10(7):1669–81. Epub 2016/02/26. doi: 10.1038/ismej.2015.235. PubMed PMID: 26905627; PubMed Central PMCID: PMCPMC4918442.

36. Friedman J, Alm EJ. Inferring correlation networks from genomic survey data. PLoS Comput Biol. 2012;8(9):e1002687. Epub 2012/10/03. doi: 10.1371/journal.pcbi.1002687. PubMed PMID: 23028285; PubMed Central PMCID: PMCPMC3447976.

37. Tett A, Huang KD, Asnicar F, Fehlner-Peach H, Pasolli E, Karcher N, et al. The Prevotella copri Complex Comprises Four Distinct Clades Underrepresented in Westernized Populations. Cell Host Microbe. 2019;26(5):666–79 e7. Epub 2019/10/15. doi: 10.1016/j.chom.2019.08.018. PubMed PMID: 31607556; PubMed Central PMCID: PMCPMC6854460.

38. Li L, Ma ZS. Testing the Neutral Theory of Biodiversity with Human Microbiome Datasets. Sci Rep. 2016;6:31448. Epub 2016/08/17. doi: 10.1038/srep31448. PubMed PMID: 27527985; PubMed Central PMCID: PMCPMC4985628.

39. Pragman AA, Knutson KA, Gould TJ, Hodgson SW, Isaacson RE, Reilly CS, et al. Chronic obstructive pulmonary disease upper airway microbiome is associated with select clinical characteristics. PLoS One. 2019;14(7):e0219962. Epub 2019/07/25. doi: 10.1371/journal.pone.0219962. PubMed PMID: 31335912; PubMed Central PMCID: PMCPMC6650084.

40. Bajorek S, Parker L, Li N, Winglee K, Weaver M, Johnson J, et al. Initial microbial community of the neonatal stomach immediately after birth. Gut Microbes. 2019;10(3):289–97. Epub 2018/11/09. doi: 10.1080/19490976.2018.1520578. PubMed PMID: 30404568; PubMed Central PMCID: PMCPMC6546338.

41. Vandeputte D, Kathagen G, D’Hoe K, Vieira-Silva S, Valles-Colomer M, Sabino J, et al. Quantitative microbiome profiling links gut community variation to microbial load. Nature. 2017;551(7681):507–11. Epub 2017/11/17. doi: 10.1038/nature24460. PubMed PMID: 29143816.

42. Fang H, Huang C, Zhao H, Deng M. gCoda: Conditional Dependence Network Inference for Compositional Data. J Comput Biol. 2017;24(7):699–708. Epub 2017/05/11. doi: 10.1089/cmb.2017.0054. PubMed PMID: 28489411; PubMed Central PMCID: PMCPMC5510714.

43. Dempster AP, Laird NM, Rubin DB. Maximum likelihood from incomplete data via the EM algorithm. Journal of the Royal Statistical Society: Series B (Methodological). 1977;39(1):1–22. doi: doi.org/10.1111/j.2517-6161.1977.tb01600.x.

44. Chacón JE, Duong T. Multivariate kernel smoothing and its applications: CRC Press; 2018.

45. Lin H, Peddada SD. Analysis of compositions of microbiomes with bias correction. Nat Commun. 2020;11(1):3514. Epub 2020/07/16. doi: 10.1038/s41467-020-17041-7. PubMed PMID: 32665548; PubMed Central PMCID: PMCPMC7360769.

46. Hosmer Jr DW, Lemeshow S, Sturdivant RX. Applied logistic regression: John Wiley & Sons; 2013.

